# Spatial transcriptomics reveals antiparasitic targets associated with essential behaviors in the human parasite *Brugia malayi*

**DOI:** 10.1101/2021.08.24.456436

**Authors:** Paul M. Airs, Kathy Vaccaro, Kendra J. Gallo, Nathalie Dinguirard, Zachary W. Heimark, Nicolas J. Wheeler, Jiaye He, Kurt R. Weiss, Nathan E. Schroeder, Jan Huisken, Mostafa Zamanian

## Abstract

Lymphatic filariasis (LF) is a chronic debilitating neglected tropical disease (NTD) caused by mosquito-transmitted nematodes that afflicts over 60 million people. Control of LF relies on routine mass drug administration with antiparasitics that clear circulating larval parasites but are ineffective against adults. The development of effective adulticides is hampered by a poor understanding of the processes and tissues driving parasite survival in the host. The adult filariae head region contains essential tissues that control parasite feeding, sensory, secretory, and reproductive behaviors, which express promising molecular substrates for the development of antifilarial drugs, vaccines, and diagnostics. We have adapted spatial transcriptomic approaches to map gene expression patterns across these prioritized but historically intractable head tissues. Spatial and tissue-resolved data reveal distinct biases in the origins of known drug targets and secreted antigens. These data were used to identify potential new drug and vaccine targets, including putative hidden antigens expressed in the alimentary canal, and to spatially associate receptor subunits belonging to druggable families. Spatial transcriptomic approaches provide a powerful resource to aid gene function inference and seed antiparasitic discovery pipelines across helminths of relevance to human and animal health.

## INTRODUCTION

Lymphatic filariasis (LF) is a chronic and debilitating neglected tropical disease (NTD) recognized as a leading global cause of long-term disability. Over 60 million people are currently infected with LF and ~900 million people are at risk of infection across 72 endemic countries.^1–3^ LF is caused by adult stage *Brugia malayi, Brugia timori*, and *Wuchereria bancrofti* filarial nematodes, which reside in the lymphatics and produce microfilariae that migrate to the blood and undergo cyclodevelopmental transmission in competent blood-feeding mosquito vectors.^4^ Adult parasites induce inflammation and blockage of lymph that can result in disfiguring and stigmatizing^5,6^ manifestations, including lymphedema (most notably elephantiasis) and hydrocele that afflicts an estimated 36 million individuals.^1,2^

LF control relies on routine mass drug administration with anthelmintics, which effectively clear microfilariae but are ineffective against adult stages and are contraindicated in areas co-endemic for other filarial parasites.^1,2,7^ Anthelmintic resistance is widespread in veterinary medicine and also represents a threat to filariasis control efforts in both animals and humans.^8–12^ To address these challenges and accelerate LF elimination there is a need to generate new antifilarial therapies, particularly drugs effective against adult stage parasites. Current anthelmintics target or dysregulate parasite cell integrity, neuromuscular control, reproductive potential, and the secretion of parasite molecules necessary for the establishment and maintenance of parasitism.^13–18^ The development of macrofilaricidal (adult-killing) drugs can be hastened by an improved knowledge of tissues that underpin survival in adult parasites.

In adult filarial parasites, vital tissues and interfaces for host-parasite communication are concentrated within the anteriormost region of the body plan. The first millimeter of the *B. malayi* female head region (~3% of the length) contains cells and tissues that control parasite feeding, sensory, secretory, and reproductive behaviors.^19^ Transcriptomic profiling of this region can aid the prioritization of new antifilarial targets, localize the targets of existing drugs, and provide clues to the origins of immunomodulatory molecules released into the host environment. This effort is currently impeded by a lack of scalable transgenesis and *in situ* localization techniques in this two-host parasite system.

Bulk transcriptomics in filarial parasites has thus far been used to explore changes in gene expression associated with development^20–23^ and environmental or drug perturbations^24–27^. While proteomics has shed light on large and accessible tissues in *B. malayi^28^*, small head-associated structures are massively underrepresented in whole-parasite omics and have yet to be characterized. Here, we adapt spatial transcriptomic and microscopy approaches to profile the head region of *B. malayi* and resolve gene expression patterns in critical tissues at the host-parasite interface. RNA tomography^29,30^ and tissue-specific transcriptomes are leveraged to map the distributions of current drug targets and known antigens, as well as to prioritize putative antiparasitic and vaccine targets. The first application of these complementary methods in a human parasitic nematode provides a template for the localization of gene transcripts and targets of therapeutic and diagnostic value in similarly intractable parasitic nematodes of human and veterinary significance.

## RESULTS

### The adult filarial head region expresses prominent antigens and known drug targets

Adult stage filariae cause incurable chronic illnesses. To develop new therapies that aid parasite elimination, we must learn more about tissues and structures underlying adult behaviors. Adult female *B. malayi* are ~34.6 mm (31.8—39.8 mm) in length when reared in Mongolian jirds, but the vast majority of their body plan is composed of mid-body structures including the body wall muscle, the reproductive tract, and intestine.^19^ The anterior-most 3% (~1 mm) of the parasite head region contains vital structures including the buccal cavity, amphid neurons, nerve ring, vulva, pharynx, esophageal-intestinal junction, and the excretory-secretory (ES) apparatus. These tissues control essential parasite behaviors and include host-parasite interfaces where drug and antigen interactions likely occur.

To identify head-enriched gene transcripts, individual adult female *B. malayi* head regions were dissected from the body at the vulva (~0.6 mm from anterior) using ultra-fine probes (**Figure 1A**). The vulva was chosen as a visible marker to ensure head tissues were captured and isolated from the reproductive tract, which would be contaminated with microfilaria. Low-input RNA-seq was carried out using paired head and body tissues isolated from three individual parasites. Biological replicates displayed high concordance (**Figure 1B**), with reads from head and body region samples uniquely mapping to the *B. malayi* genome at 70-80%. Analysis of differentially-expressed genes (DEGs) identified 2,406 head-enriched genes (log_2_(FC) > 1 and p-value < 0.01) with at least 30 total raw reads from the six samples (**Supplementary Table 1**). Transcripts associated with secreted proteins^31,32^ are distributed evenly across both head and body region tissues, suggesting mixed origins for what are classically referred to as “ES products” (**Figure 1C**). In striking contrast, nearly all (86%) prominent or abundant filarial antigens with known immunomodulatory capacity, including those that have been pursued as vaccine candidates^33^, are head-enriched (**Figure 1C**).

**Figure 1:**
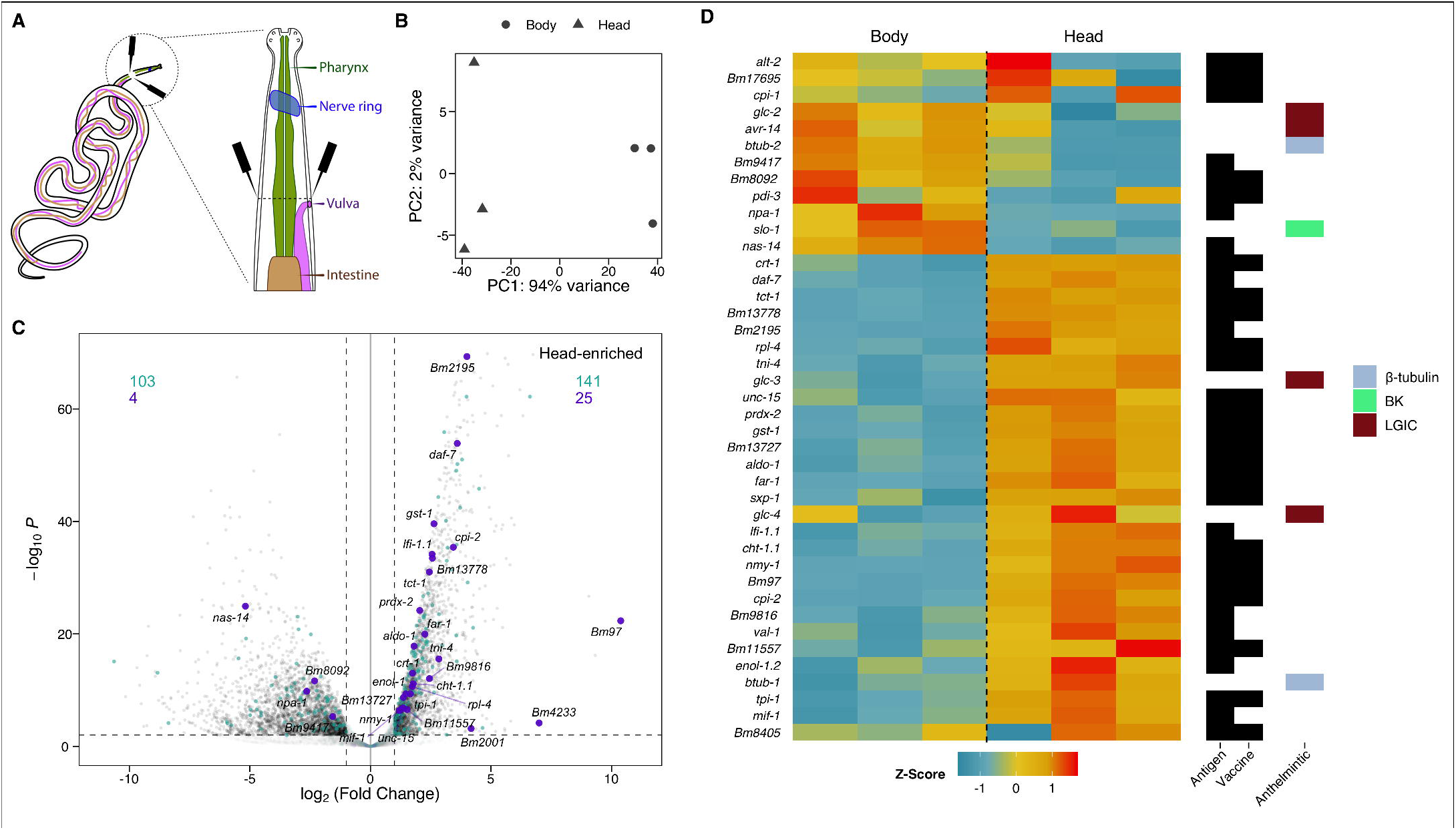
The *B. malayi* head transcriptome reveals enrichment of anthelmintic targets, prominent antigens, and vaccine candidates. (A) Illustration of head region tissues and point of dissection adjacent to the vulva (black probes). (B) Principal component analysis (PCA) plot of head and corresponding body transcriptome replicates. Each point represents tissue from a different parasite. (C) Volcano plot highlighting head enrichment of prominent antigens and immunomodulatory genes (purple)^16,33,94–101^, compared to a more diffuse pattern of expression for proteins identified in secretome studies (teal)^31,32^. (D) Heatmap of gene expression patterns shows head-enrichment of candidate vaccine targets, as well as both head- and body-enrichment for the targets of antifilarial drugs (benzimidazoles: β-tubulins, macrocyclic lactones: GluCls, and emodepside: BK channel).

Antifilarial targets from existing classes of drugs show different distributions (**Figure 1D**). The putative glutamate-gated chloride channel (GluCl) targets of ivermectin, *Bma-avr-14* and *Bma-glc-2^34^*, show higher relative expression in body tissues consistent with *Bma-avr-14* localization to the reproductive tract and developing embryos.^35^ *Bma-glc-3* and *Bma-glc-4* channel subunits are more enriched in the head and may also play a role in macrocyclic lactone responses. *Bma-slo-1*, a putative target of emodepside, an emerging candidate adulticide for treatment of river blindness^36^, is more highly expressed in the body. Conversely, the likely β-tubulin target of albendazole (*Bma-btub-1*), based on homology to *Caenorhabditis elegans ben-1*, is head-enriched.

### Microscopic investigation of the adult head region and putative excretory-secretory apparatus

Organizational knowledge of prominent head structures can scaffold eventual spatial transcriptomic data. The buccal cavity (~5 μm), vulva (~657-667μm) and esophageal-intestinal junction (~861-1010 μm) are precisely mapped in adult female *B. malayi^19^*, but identifying the excretory-secretory (ES) system in adult stage *B. malayi* and other filaria has been notoriously difficult.^37–41^ In microfilariae, the ES apparatus is a hallmark and essential structure consisting of a pore and vesicle leading to a single excretory cell via a cytoplasmic bridge.^42^ Ivermectin is thought to disrupt microfilarial ES protein and exosome release through binding to ion channels in the vicinity of the ES vesicle^14,16,17^; however, these structures become inconspicuous through development.^37,38,40,43,44^ To help pinpoint the ES in adult female *B. malayi*, the relative organization of head structures across Clade III^45,46^ nematodes was collected from available literature (**Figure 2A**). Among Clade III parasites, the ES pore and/or cell are located posterior to the nerve ring in 23/24 species, anterior to reproductive openings (24/24 species), and anterior to the esophageal-intestinal junction (23/24 species) in at least one life stage surveyed. The conservation of structural organisation across developmental and evolutionary time indicates the presence of ES structures between the nerve ring and vulva in adult female *B. malayi*.

**Figure 2:**
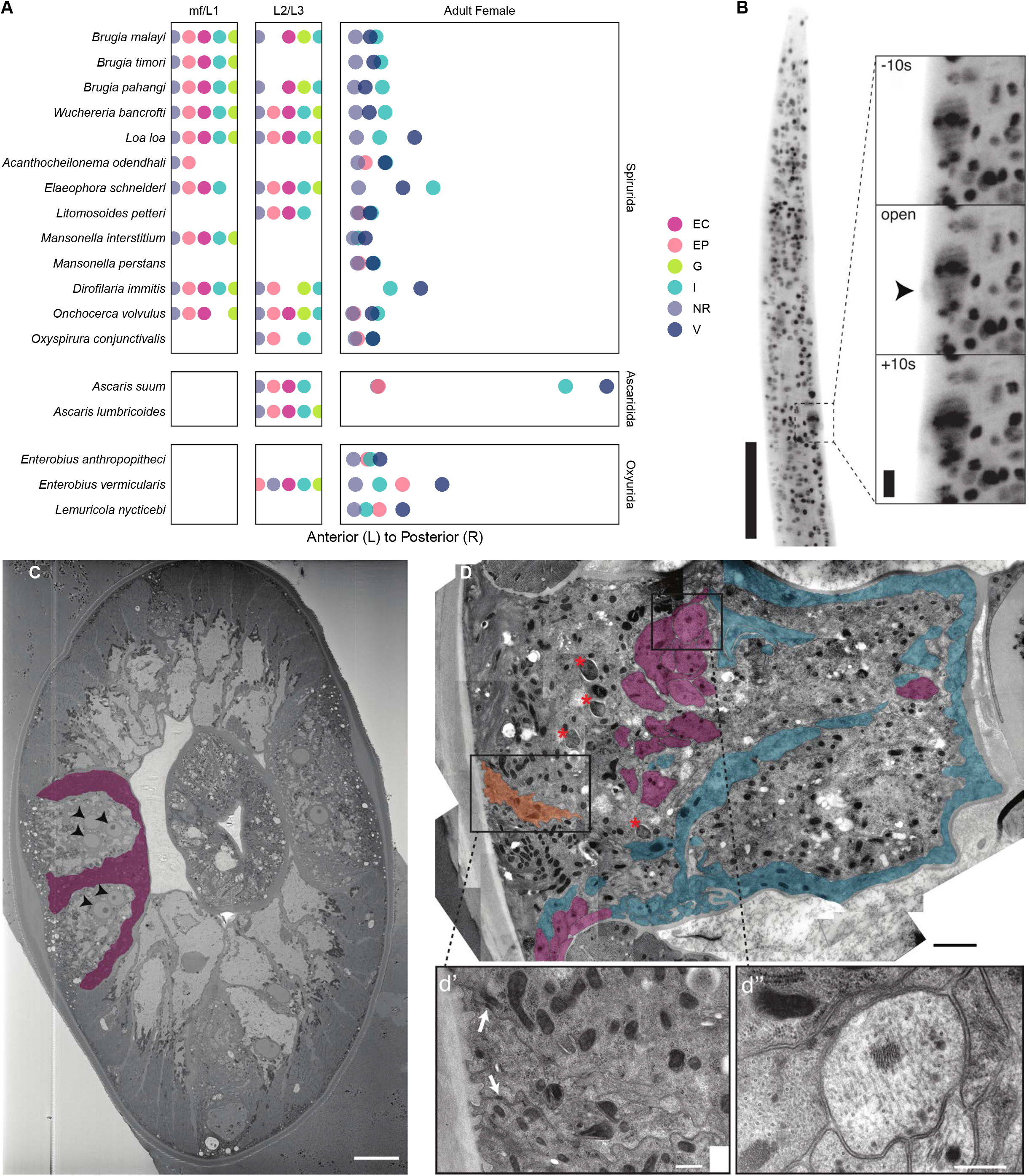
Coordinating the elusive excretory-secretory system in the adult *B. malayi* head region. (A) Comparative anatomy of Clade III nematode head structures from published descriptions (detailed in **Supplementary File 1**). Positions of the nerve ring = NR, excretory-secretory pore and/or vesicle = EP, excretory cell = EC, genital primordium of larvae = G (larvae), vulva = V, and esophageal-intestinal junction = I shown as rank order for larvae and average micron distances from anterior in adult stages. (B) Light sheet maximum intensity projection of ES pore pulsing activity (arrowhead) in DRAQ5 stained live adult male *B. malayi*. Scale bars = 100 μm and 10 μm for insets. (C) Single section from adult female SBF-SEM showing multinucleated (arrowheads) epidermis within the lateral cord and membranous structures (pseudocolor purple) that are embedded within and surround the lateral cord. Scale bar = 10 μm. (D) TEM of lateral cord in adult male showing likely seam cell homolog (pseudocolor orange), membranous processes enriched in microtubules (pseudocolor purple), membranous processes lacking obvious microtubules (pseudocolor blue), and *Wolbachia* endosymbionts (asterisks). (d’) Closeup of seam cell, identifiable by the position on the median ridge of the lateral cord and the presence of adherens junctions (arrows) connecting to surrounding epidermis. Scale bar = 2 μm. (d”) Closeup of membranous process enriched in microtubules and surrounded by epidermis. Scale bar = 400 nm.

To identify the ES pore in adults we optimized live 4D light sheet microscopy as well as multiple electron microscopy methods. Critical point drying scanning electron microscopy (SEM) of adult *Brugia* and the closely-related but much larger filarial parasite *Dirofilaria immitis* allowed clear visualization of the vulva, but not the ES pore (**Supplementary Figure 1**). This may be due to the small size and angle of the pore opening.*^47^* Light sheet imaging was adapted for live adult males partially paralyzed with 1 mM levamisole to restrict gross muscle movement, and adults were monitored for up to 1 hr at 10 s intervals. Males were chosen to avoid confusion with the confounding activity of the vulva, which is proximal in females. Nuclei stained head regions revealed instances of pulsing during which stain condensed into a large cell with a pore-like tubular structure that was then cleared from the worm ~430 μm from the anterior (**Figure 2B**, **Supplementary Video 1**). This location is consistent with the ES pore location (397-537 μm) in the fur seal parasite *Acanthocheilonema odendhali*^48^, the only filarial nematode where the adult stage ES pore has been morphometrically characterized. To our knowledge, these pulses represent the first evidence of dynamic ES pore opening events in a mammalian parasitic nematode.

To obtain a finer description of head structures and potential ES channels, we utilized high-pressure freeze fixation with serial block face-SEM (SBF-SEM) to obtain approximately ~1000 serial sections (~70 nm/section) from the anterior of an adult female (**Figure 2C**, **Supplementary Video 2**). The ventral nerve cord and pharynx were present throughout and we observed 30 pharyngeal, 21 body wall muscle (~5 per muscle quadrant), 5 ventral nerve cord, and 83 lateral cord (~40 per cord) nuclei. Similar to *C. elegans*, the lateral cords appear to be partially composed of epidermal syncytia with multiple closely apposed nuclei (**Figure 2C**). We did not observe any nuclei within the dorsal cord. The *C. elegans* excretory canal^49^ is visible in EM sections immediately ventral of the lateral cords, while in *Onchocerca volvulus* a glomerulus like excretory-structure^50^ is suggested to be embedded within the lateral cords. Neither canal-type was observed in our SBF-SEM data. Their absence is possibly due to individual variation in the position of ES structures or the excretory system may be greatly reduced in size, as proposed previously.^38,50^

Within lateral cords we observed membrane bound processes along the lateral and basal edges. These processes were described previously as axons or infolded membranes.^41,51,52^ In some regions processes are also embedded within the lateral cord, while others appear to bisect the lateral cord. Similar processes embedded in the lateral cord are not seen in *C. elegans.^53^* To better define the lateral cord in this region we turned to transmission electron microscopy (TEM) (**Figure 2D**). As previously suggested, membrane processes embedded within the lateral cord appear to be neuronal, as evidenced by numerous microtubules. However the processes located on the basal boundary of the lateral cord lacked microtubules. The absence of consistent microtubules argues against a solely commissure identity. Another possibility is that these structures towards the interior of the worm comprise a modified excretory system. Their position adjacent to the pseudocoelom would be consistent with an excretory system; however, additional serial data are needed to identify the nature of these structures.

TEM also demonstrated the presence of likely homologs to the *C. elegans* seam cells along the median ridge surrounded by putative epidermal syncytia (**Figure 2D**). These are readily identifiable by their position and the presence of adherens junctions connecting the seam to the syncytial epidermis. In *C. elegans* and other nematodes, the seam cells have stem cell-like properties and act to contribute nuclei to the growing epidermal syncytia.^54,55^ As previously shown, the lateral cords were also enriched in *Wolbachia* endosymbionts.^52^

### Spatial transcriptomics maps antiparasitic targets associated with essential tissues

To deconvolute gene expression patterns across the adult female *B. malayi* head-region we adapted and optimized RNA tomography from model organisms.^29,30^ Individual adult females were oriented and cryo-embedded (**Supplementary Video 3**) for collection of 20 μm sections along the anterior-posterior axis. Cryosection imaging was used to validate the tissue collection protocol (**Figure 3A**) and generate estimates of nuclei density at 20 μm resolution across the targeted region (**Figure 3B**). Each cryosection contains a mean of 10.15 (±1.46) nucleated cells anterior to the vulva, with decreased cell densities at the anterior tip and prior to the appearance of the ovarian tract. These observations correspond to live light sheet imaging data, with approximately 9.70 and 9.24 nuclei per 20 μm in females and males, respectively (**Figure 3C**).

**Figure 3:**
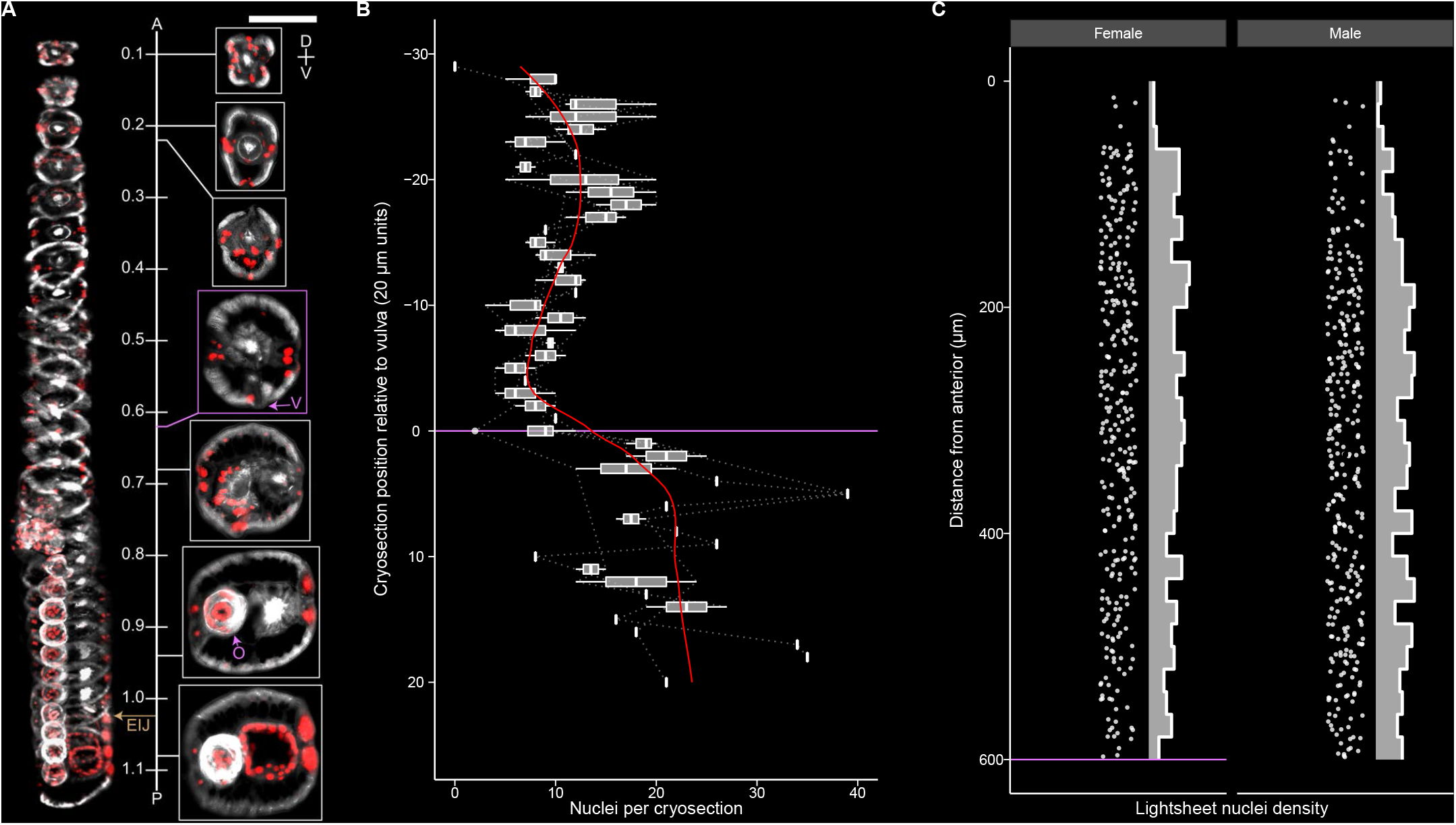
Optimization of spatial cryo-embedding and nuclei distribution in adult *B. malayi*. (A) The integrity and position of tissues is not impacted by cryopreservation and is accurately captured for RNA tomography. 3D rendered (left) and representative (right) views of 20 μm cryosections covering the anterior most portion of the adult female head highlighting the vulva = purple V, ovarian tract = purple O, and esophageal-intestinal junction = brown EIJ. DAPI = red, phalloidin = grey, A = anterior, P = posterior, D = dorsal, V = ventral, scale bar = 100 μm, numbers are mm from anterior. (B) Adult female nuclei counts per 20 μm cryosection relative to the vulva and start of the ovarian tract (purple line). (C) Adult male and female DRAQ5 stained nuclei density, captured anterior to the vulva (purple line) with light sheet imaging.

Single-worm RNA tomography was performed via sequential capture and 96-well plate based processing of individual 20 μm sections for low-input RNA-seq (**Figure 4A**). Read mapping rates are negligible (< 1%) through the first 11 sections, reflecting a conservative capture strategy to avoid missing the anterior tip of the head, and rise to 70% (± 14%) through the remaining sections. Hierarchical clustering of three RNA tomography replicates reveals unique gene expression signatures across sections (**Figure 4B and C**). Robust genes were defined as those associated with >20 counts across all sections for a given replicate and >10 counts in a single section. Robust sections were defined as those expressing at least 100 genes with >10 counts. 8,900 genes are detected across the head of the first (highest-quality) replicate (**Figure 4D**). Additional replicates exhibited lower gene coverage, but shared 97% and 94% of their detected genes with the first run (**Figure 4E**). 5,810 genes were detected across all replicates, including 2,375 of 2,406 (98.7%) transcripts previously identified as head-enriched. RNA tomography captured approximately 48% of *B. malayi* protein-coding genes, reflecting the great diversity of tissue and cell types contained within the relatively small head region.

**Figure 4:**
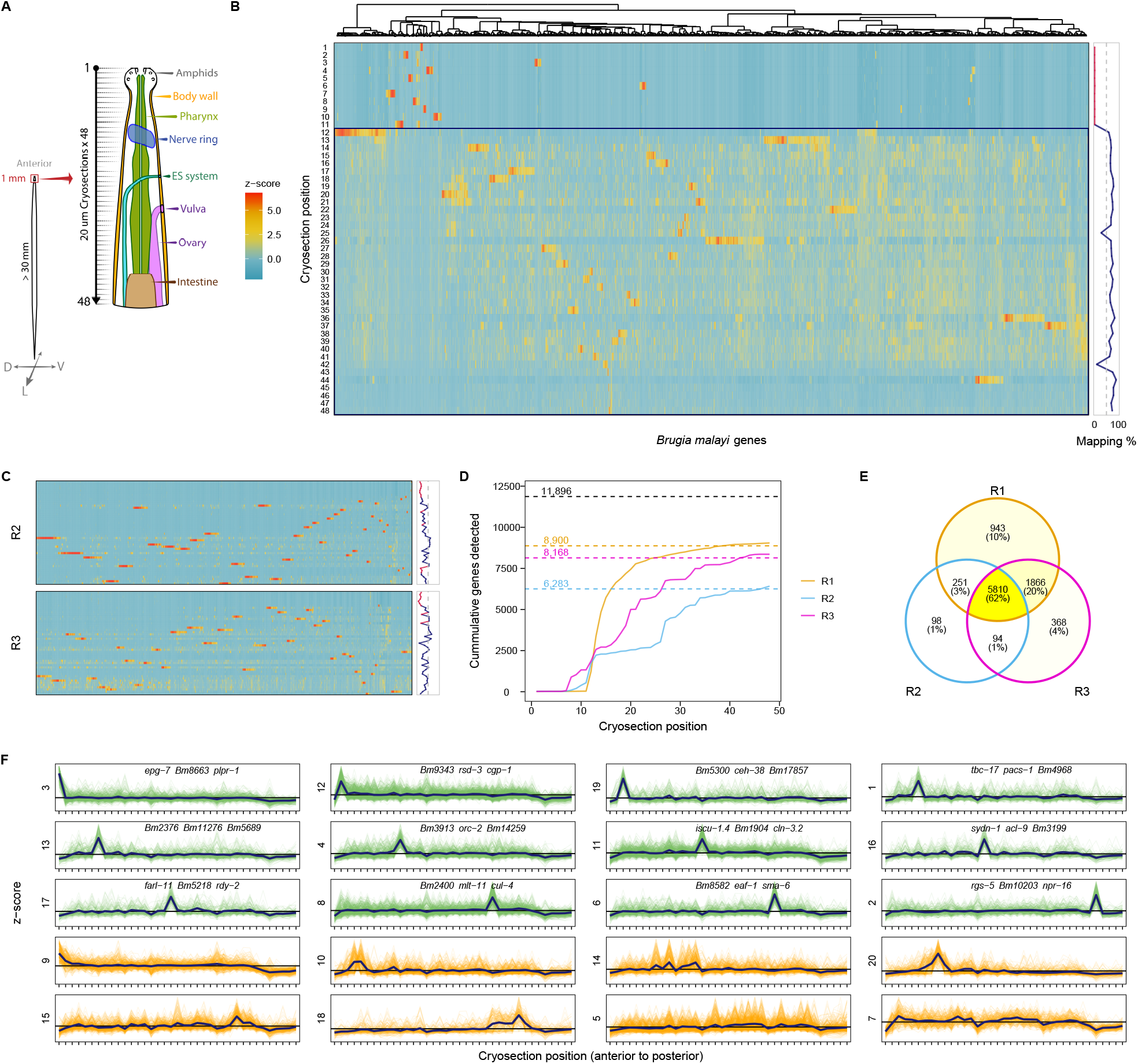
RNA tomography of the *B. malayi* adult female head. (A) Graphical representation of target tissue organization in the adult female *B. malayi* head investigated with RNA tomography. (B) Spatial gene expression heatmap and read-mapping rates for *B. malayi* RNA tomography replicate 1. High-quality cryosections associated with high rates of uniquely mapped reads fall below the blue line (cryosection 12). Z-scores reflect scale-normalized TPM counts. (C) Spatial gene expression heatmaps for additional replicates (R2 and R3). (D) Cumulative protein-coding genes identified along the anterior-posterior axis for each replicate. Genes were included in the count if they were found in at least one slice with > 10 raw reads. (E) Overlap of robustly-expressed genes detected across replicates. (F) Clustering of spatial expression patterns from the highest quality RNA tomography run (x-axis: crysections 12-48). Cluster IDs (**Supplementary Table 2**) are shown for both localized (green) and diffuse (orange) spatial expression patterns. The three highest expressed genes are provided as markers for peaks localized to a single cryosection.

Genes were grouped by spatial expression pattern, displaying either localized or diffuse expression patterns down the anterior-posterior axis. The former likely represent gene transcripts and markers restricted to distinct neurons, while the latter reflect recurring cell types such as the epidermis, body wall muscle, or pharynx (**Figure 4F** and **Supplementary Table 2**). Prominent secreted antigens (e.g., *Bma-mif-1, Bma-tpi-1, Bma-cpi-2*), including proteins associated with exosomes (e.g., *Bma-lec-1* and *Bma-enol-1*)^17^, do not fall into a specific cluster, reinforcing the heterogeneous nature of their transcriptional origins even if ultimately released from the same orifice. Neighboring cryosections were coalesced to map the most abundantly expressed genes with respect to head structures. The most highly expressed pre-vulval genes are the neuropeptide-like protein *Bma-nlp-77* and the collagen *Bma-col-72*, replaced by the immunogens *Bma-val-1^56^* and *Bm97^57^* where the vulva is expected to appear (**Figure 5A**). A fraction of post-vulval transcripts are likely associated with progeny in the reproductive tract.

**Figure 5:**
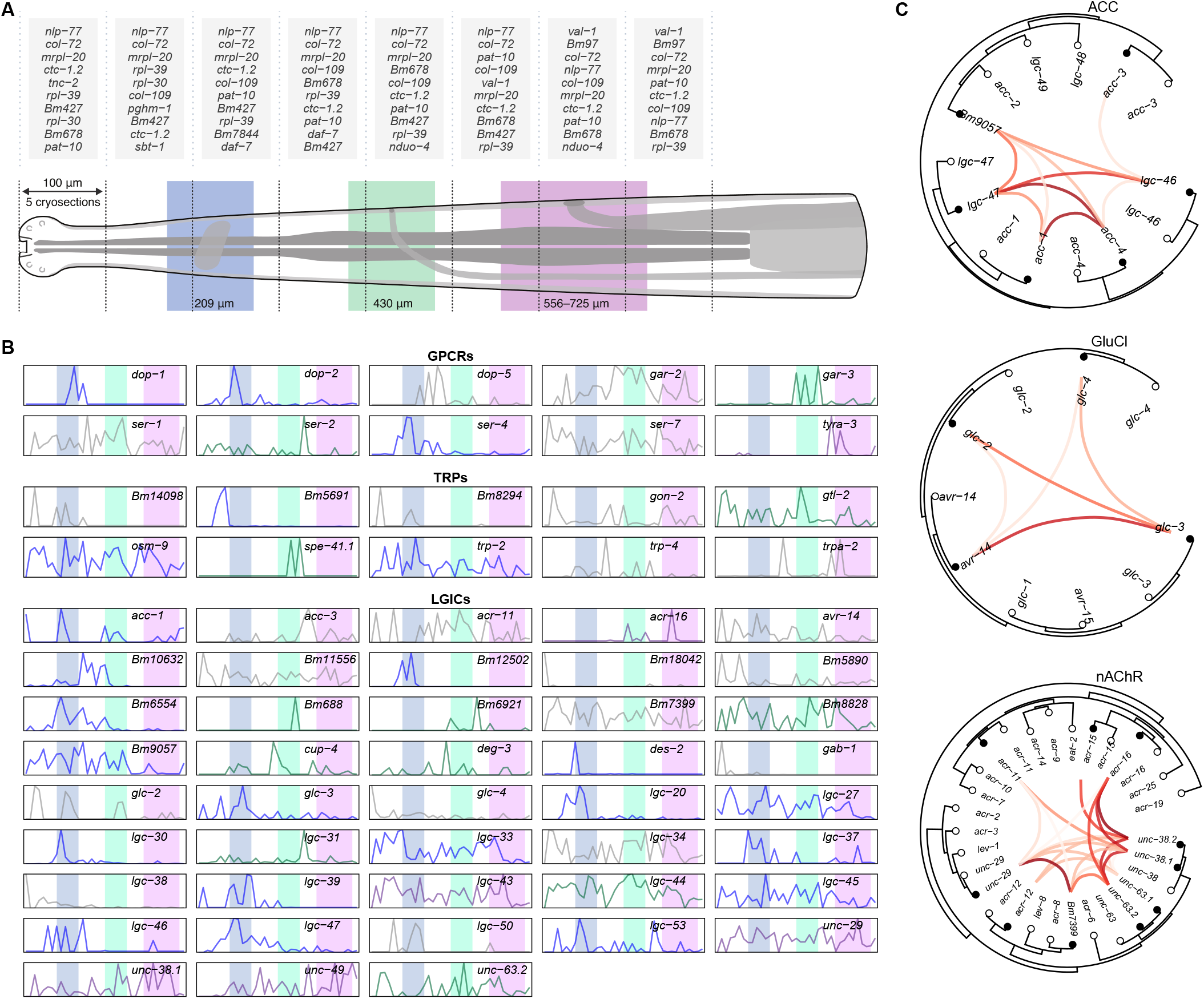
Spatial analyses of highly-expressed genes and putative receptor drug targets. (A) The most-highly expressed genes are shown for each 100 μm pseudo-section (combining neighboring sets of five 20 μm cryosections), in relation to the anatomy of the adult female head. The location ranges of major structures reflect worm-to-worm variation: ± 50 μm for the nerve ring (blue) and ES pore (green), and as empirically determined for the vulva (purple). (B) Spatial patterns of expression for detected genes (TPM > 10 in at least one cryosection) belonging to classically druggable receptor families (GPCRs, TRPs, LGICs) for the highest-quality biological replicate (y-axis = scaled TPM). Line colors signify transcripts that are largely restricted to or are enriched in the vicinity of the matching head structure. (C) Phylogenetic trees showing spatial correlations among cys-loop LGIC subunits belonging to the acetylcholine-gated chloride channel (ACCs, top), glutamate-gated chloride channel (GluCls, middle), and nicotinic acetylcholine receptor (nAChRs, bottom) subfamilies. Links represent the Pearson correlation coefficient, which was calculated from scale-normalized TPM values from the highest-quality RNA tomography replicate; only links with a positive correlation are shown (black points: *B. malayi* and white points: *C. elegans*).

We next examined the spatial distributions of druggable receptor and ion channel families, detecting 10 (of 11) transient receptor potential (TRP) channel subunits, 43 (of 52) cys-loop ligand-gated ion channel (LGIC) subunits, and 10 (of 11) aminergic G protein-coupled receptors (GPCRs) across the head region (TPM > 10 in at least one cryosection). A subset of these receptors are restricted to or enriched in the vicinity of the nerve ring, ES pore, or vulva (**Figure 5B**), suggesting an outsized role in neuromuscular control of movement, secretion, or fecundity. LGICs represent the most successful class of anthelmintic targets, mediating the antiparasitic effects of nicotinic receptor agonists and macrocyclic lactones. These pentameric channels can be pharmacologically characterized in heterologous cells^58^ but it is unknown whether heteromeric channels functionally constituted in surrogate systems reflect endogenous channel subunit interactions. To guide heterologous studies, we used spatial correlations among channel subunits for major LGIC subfamilies (**Supplementary Figure 2**) to predict subunits that are more likely to be found in the same cells and form functional channels (**Figure 5C**).

### Discovery of candidate hidden antigens in the pharynx and intestine

The nematode pharynx and gut are established target sites for existing^59–61^ and emerging anthelmintics^62^. The alimentary tract is also a critical host-parasite interface and potential sources of ‘hidden’ antigens for vaccine development.^28,63,64^ These antigens may evade host immune recognition but remain accessible to vaccine-induced antibodies in the course of parasite feeding^64^ - the rationale behind the protective immunity offered by the first commercial parasitic nematode vaccine in ruminants^65^. While RNA tomography provides an anterior-posterior map of gene expression across the head region, these data alone cannot be used to cleanly infer pharyngeal or intestinal transcripts.

To separately capture these tissues, whole intestines were first isolated by microdissection of live adult females (**Figure 6A**), revealing 1,077 intestine-enriched genes (log_2_(FC) > 1 and p-value < 0.01) with 489 genes predicted to contain at least one transmembrane domain. 64 putative membrane proteins were further prioritized as candidate hidden antigens based on high intestinal expression (mean TPM > 100) and relatively low abundance in non-intestinal tissues (intestinal:non-intestinal TPM ratio > 10) (**Figure 6B** and **Supplementary Table 3**). These data greatly expand on the *Brugia* intestinal proteome^28^ and provide new leads that are more likely to be tissue restricted. We identify cathepsin-like protease *Bma-cpl-1* as an intestinally-enriched target, along with membrane targets that include a GABA receptor subunit (*Bma-gab-1*), glutamate transporter (*Bma-gtl-1*), neuropeptide GPCR (*Bma-npr-23*), genes associated with hypersensitivity to pore-forming toxins (*Bma-hpo-8* and *Bma-hpo-28*), and an ortholog of a *C. elegans* 7TM intestinal receptor involved in innate immune responses (*Bma-fshr-1*) (**Figure 6C**).^66^

**Figure 6:**
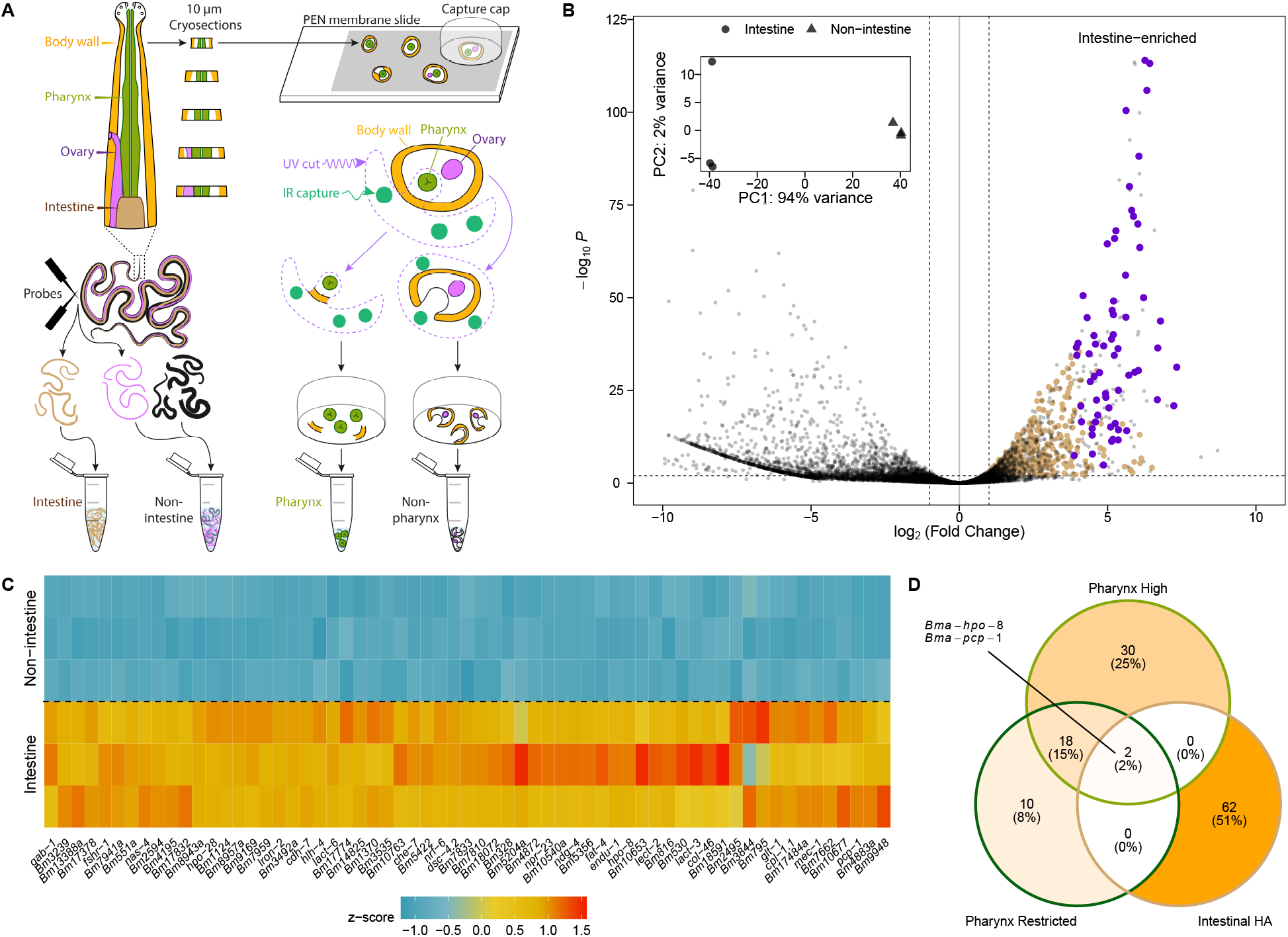
Alimentary-specific transcriptomics of *B. malayi* adult female pharynx and intestine. (A) Graphical representation of dissection and laser capture microdissection strategies to recover tissues of the lower (intestine) and upper (pharynx) alimentary canal. (B) Volcano plot highlighting intestinally-enriched transcripts from paired intestinal and non-intestinal samples of three biological replicates (individual parasites). Transmembrane proteins are considered candidate hidden antigens if they are intestine-enriched (brown), with a prioritized subset that is intestine-restricted (purple). (C) Expression heatmap of prioritized intestine-restricted hidden antigens. z-scores were generated from vst-transformed counts. (D) Overlap of candidate intestinal hidden antigens, highly-expressed pharyngeal genes (top 50 mean pharyngeal TPM among genes with > 100 TPM for each of three replicates), and pharyngeal genes with more tissue-restricted profiles.

To determine whether targets in the intestine were conserved in the upper alimentary tract we performed laser capture microdissection (LCM) of pharyngeal tissue. Pharyngeal and adjacent non-pharyngeal tissue were isolated by LCM from 10 μm thick adult female head sections generated by the RNA tomography cryosectioning technique (**Figure 6A**). Each collected sample, estimated to contain fewer than five cells, was subject to low-input RNA-seq. Pharyngeal samples cluster as expected, while non-pharyngeal samples are divergent, reflecting collections of disparate cell types from different positions in the head (**Supplementary Figure 3**).

Focusing on transmembrane proteins, we observe little overlap between the most highly expressed genes in the pharynx, including those more restricted to the pharynx (pharyngeal:non-pharyngeal TPM ratio > 10), and prioritized intestinal targets (**Figure 6D** and **Supplementary Table 4**). This suggests that the upper and lower alimentary canal are sources of unique targets and candidate hidden antigens. *Bma-hpo-8* and *Bma-pcp-1*, a membrane-bound peptidase, represent exceptions enriched across the alimentary canal. Both intestinal and pharyngeal hidden antigen candidates are comprised mostly of genes of unknown function and many are orthologous to extra-intestinal *C. elegans* genes. This highlights the need for care in ascribing functions and spatially mapping tissue-specific markers from this clade V model nematode to distantly-related clade III parasites.

## DISCUSSION

Spatially resolved gene expression patterns and tissue-specific transcriptomes can aid our functional understanding of genes^67^, especially in non-model organisms where transgenesis and functional genomics are not routine. To this end, we have generated the first genome-wide expression maps at fine scale in a multicellular parasite, focusing on the *B. malayi* adult head region. This tightly organized space encompasses tissues and structures responsible for vital sensory, secretory, reproductive, and feeding processes required for parasite survival and the maintenance of infection in the human host. We adapted low-input tissue capture and RNA tomography, combined with light-sheet and electron microscopy, to survey anterior-posterior expression patterns and map these data to tissues of interest.

Openings in the *B. malayi* head act as druggable host-parasite interfaces and as potential conduits for prominent secretory antigens^31,68^ and vaccine candidates^28,33,65^. Secretomes of adult stage *B. malayi* are well defined^13,31^ but the anatomical origins of these products are unknown, owing in part to difficulty identifying the adult ES-pore.^37,38,40,41,43,44^ We show that the great majority of prominent antigens are head-enriched, but do not fall into a specific spatial pattern within the head region. Complementary light sheet and electron microscopy efforts provide the first direct evidence of ES pore activity in a mammalian parasitic nematode and suggest a potentially contracted or modified ES system in the adult stage^38,50^, which requires further microscopy-based investigation.

Spatial and tissue-specific transcriptomics data were leveraged to map the distributions of current anthelmintic targets and to prioritize new drug and vaccine targets, including candidate membrane-anchored ‘hidden’ antigens that are highly-expressed and more likely to be restricted to the upper or lower alimentary canal. Transcripts encoding for proteins that belong to traditionally druggable receptor families were associated with the estimated locations of the nerve ring, ES pore, and vulva. These receptors may serve as targets for the dysregulation or inhibition of parasite neuromuscular control, host-parasite communication, and fecundity, respectively.

While these spatial transcriptomics techniques provide positional information in the context of a whole organism or region, they do not capture individual cells and are challenging to align given the significant anatomical and size variation observed among adult stage filarial parasites. To complement RNA tomography, single-cell approaches in parasitic nematodes, such as those applied in *C. elegans^69^*, can provide more granular information about cell and tissue-specific expression patterns. However, unlike *C. elegans*, there are no validated tissue-specific markers to map cell lineages in *B. malayi*. Transgenic approaches are developing^70^, but cannot conceivably be scaled given the challenges of the two-host life cycle. Ultimately, the combination of spatial and single-cell approaches in *B. malayi* provides a pathway to marry transcription to specific cells within defined tissues of interest.

Taken together, these findings highlight the utility of spatial transcriptomic techniques when applied in parasitic nematodes and show that the resulting data can be used to define region and tissue specific gene expression patterns in small and densely packed parasite tissues.

## METHODS

### Parasite culture

*Brugia malayi* adults (NIH-FR3) extracted from the *Meriones unguiculatus* infection system were maintained in daily changes of RPMI 1640 with L-glutamine (Sigma-Aldrich) supplemented with FBS (10 % v/v, Fisher Scientific) and penicillin-streptomycin (100 U/mL, Gibco) at 37°C with 5% CO_2_ unless otherwise specified. Individual adults were separated by sex into 3-4 mL of culture media. For RNA-seq analyses, individual worms were acclimated in culture for 18-24 hrs prior to fixation, preservation, or RNA extraction unless otherwise stated.

### Parasite tissue dissections

A modification of the Morris *et. al.^28^* method was employed where individual adult female *B. malayi* were washed 3x in nuclease-free PBS then dissected using Eliminase (VWR) cleaned 0.15 mm minuten pin dissecting probes (Bioquip) in PBS in a petri dish under a ZEISS Stemi 508 with Sony Exmor CMOS IMX178 camera. For head vs body RNA-seq, the head was severed by crossing two probes adjacent to the vulva. For the intestine vs carcass RNA-seq, the body was held in place using one probe and another was used to gently nick and pierce the cuticle at the midbody, releasing internal organs. The head and tail were then severed to free intestinal tract ends. Eliminase washed No. 5 forceps were used to pull the intestine away from the body. Individual tissues were transferred to 2 mL Safe-Lock tubes (Eppendorf) containing 300 μL TRIzol LS (Invitrogen) and 100 μL nuclease-free water, vortexed, flash frozen in liquid nitrogen, and stored at −80°C. For RNA extraction, samples were thawed and homogenized with a single 3 mm Eliminase washed stainless steel ball bearing in for 3 min at 30 Hz (TissueLyser II, Qiagen) then purified by the Direct-zol RNA microprep kit (Zymo).

### RNA tomography

Individual adult female *B. malayi* were washed thrice in RPMI 1640 with L-glutamine, soaked in RPMI 1640 with 0.005% methylene blue for 2 min, then washed once more with RPMI 1640. Stained worms were picked using Eliminase cleaned 0.15 mm minuten pin dissecting probes directly into clear TissueTek O.C.T. (Electron Microscopy Sciences) in a Stainless Steel Base Mold (Simport Scientific). The mold was positioned immediately prior to staining on a bed of dry ice under a ZEISS Stemi 508 with Sony Exmor CMOS IMX178 camera. Once in O.C.T., the body was straightened so that the head region was positioned parallel to the long face of the mold using the probe. A ~0.5 μL methylene blue (0.1% in water) dye dot was then placed roughly 1 mm above the anterior-most portion of the head to act as a location marker. During freezing the position of the worm was imaged in relation to the edge of the mold and the dye marker to calculate distance between the block edge and the dye dot as well as the dye dot to the sample. Frozen O.C.T. blocks were covered in parafilm, indexed, and stored at −80°C until sectioning. Cryosections (48 x 20 μm) along the anterior-posterior axis were taken on a Leica CM3050 S Research Cryostat by positioning O.C.T. blocks perpendicular to the cutting face. The number of sections required to pass through the dye dot was used to estimate the number of sections required to reach the worm head based on images taken during embedding. A 40x loupe was used to interrogate the O.C.T block for signs of the methylene blue stained sample. Edges of the O.C.T block were shaved to minimize O.C.T. contamination and 20 μm sections were kept individually on dry ice during sectioning. Sections were thawed in 75 μL nuclease-free water for 30 s, pipette mixed with 225 μL TRIzol LS, and purified with the Direct-zol 96 kit (Zymo, eluted in 20 μL of nuclease-free water).

### Laser-capture microdissection (LCM) of the adult female pharynx

Sections (10 μm) were collected using the RNA tomography sectioning protocol and placed directly onto UV irradiated (254 nm, Stratalinker) PEN Membrane Glass Slides (Applied Biosystems). Slides were rinsed once in nuclease-free water for 1 min to remove O.C.T, followed by a 1 mL wash series in 70%, 90%, 95%, and 100% ethanol, then air dried for 10 min and stored at −80°C in a 50 mL conical tube. Slides were equilibrated to room temperature before being loaded onto an ArcturusXT LCM Instrument. Sections were inspected for the presence of a pharynx, and captured under CapSure HS LCM Caps (Applied Biosystems) by UV laser (settings include UV current: 15, cutting speed: 300, pulse frequency: 500, section thickness: 10, cut: 10, tab length: 1) and captured by IR (settings include IR spots: 3, spacing: 60, diameter: 50-75, power: 99, duration: 49). Cuts were made to separate the pharynx from the rest of the tissue, leaving a tab of tissue that connects to the rest of the PEN membrane. Each cut section was collected on one CapsSure cap with the remaining tissue collected on another cap in GeneAmp Thin-Walled Reaction Tubes (Applied Biosystems), transferred to dry ice for immediate processing after collection. Three replicates were performed with 2-4 sections per replicate per group (pharynx vs non-pharynx).

### RNA-seq library preparation and sequencing

For NEBNext library preps, DNA quantity was checked by Qubit (dsDNA HS Assay Kit, Invitrogen) and SPRIselect beads (Beckman Coulter) were used for DNA purification steps. For cDNA amplification and PCR enrichment of the adapter ligated DNA, cycle numbers were optimized for each sample unless stated otherwise.

#### Head vs body

The Clontech SMARTSeq v4 Ultra-Low Input RNA kit (Takara) was used with 1.7 ng of input RNA from each sample, as determined by Agilent RNA 6000 Pico Kit on a 2100 Bioanalyzer (Agilent). Full length cDNA was quantified by 2100 Bioanalyzer. 150 pg of amplified cDNA was tagmented and index-amplified using Nextera XT adapters (Illumina). DNA quantity was assessed by Qubit (dsDNA HS Assay Kit) and quality by 2100 Bioanalyzer. Libraries were balanced by Illumina MiSeq Nano for a single lane of 1×100 bp sequencing on the Illumina HiSeq 2500.

#### RNA tomography

The first library was prepared as described for head vs body RNAseq. For additional replicates, 8 μL of RNA was added per section to step 2.1 through to 2.11.11 of the NEBNext Single Cell/Low Input RNA Library Prep Kit for Illumina (NEB) using NEBNext Multiplex Oligos for Illumina (Index Primers Set 1, NEB) and sequenced on an Illumina Novaseq 6000 (2×150 bp, S1 flow cell).

#### Intestine dissection and pharynx LCM

Libraries were prepared as previously described with the NEBNext kit. For intestine and carcass tissues, 8 μl of purified RNA was added per sample to step 2.3 through to 2.11.11. For pharynx and non-pharynx RNA, tissues on CapSure caps were transferred by Eliminase cleaned No. 5 forceps directly to 8 μl of 1x NEBNext Cell LysisBuffer, frozen at −80°C overnight for one cycle of freeze-cracking to release RNA. Lysate was submitted to step 1.3 through to step 1.12.11. For pharynx and non-pharynx RNA, 20 cycles of cDNA amplification were performed at step 1.5 and 12 cycles of PCR enrichment of the adapter ligated DNA were performed at step 1.11. Libraries were sequenced on an Illumina Novaseq 6000 (2×150 bp, 4 million reads per sample).

### Epifluorecent imaging of representative *B. malayi* cryosections

Sections (20 μm) were collected using the RNA tomography sectioning protocol, placed sequentially in rows on charged slides (Thermo Scientific) and allowed to dry at room temperature for 5 min. Slides were rehydrated in PBS for 30 s followed by marking of sections with diamond pen, rinsed in molecular grade water and air dried again. Sections were then fixed in 3.4% formaldehyde at room temperature for 15 min, washed twice with PBS and stored in 70% ethanol. Slides were stained in 20 mL of 70% ethanol containing 1 μg DAPI (Invitrogen) and 5 μL AlexaFluor 488 Phalloidin (Invitrogen) for 8 min at room temperature, washed in 70% ethanol, then rehydrated and mounted in PBS prior to imaging on a Zeiss Axio Scope A1. 3D rendering of color-merged sections was performed in Fiji^72^ using TrakEM2^73^ to orient all sections per individual as a stack, which was then compiled in 3D viewer^74^ (voxel depth 124 / 20 μm section, resampling rate = 1).

### Serial block face-SEM (SBF-SEM) and TEM

Adult female *B. malayi* were prefixed in 2% paraformaldehyde and cut with a scalpel posterior of the vulva. The anterior portion was immediately placed into a HPF 3 mm specimen carrier with 20% bovine serum albumin (Sigma-Aldrich) and frozen in an Alba HPM 010. Specimens were transferred into an RMC FS-8500 for freeze substitution. Specimens prepared for SBF-SEM were freeze substituted in 2% OsO_4_, 0.1% uranyl acetate and 2% H_2_O followed by *en bloc* Osmium-Thiocarbohydrazide-Osmium staining^75,76^ and embedding in Durcupan ACM (Electron Microscopy Sciences). Serial block-face imaging was conducted in a Zeiss Sigma 3View system with variable pressure with 30 nm x 70 nm resolution at 5.5 kV. Specimens for TEM were freeze substituted in 2% OsO_4_, 0.1% uranyl acetate, and 2% H_2_O and embedded in Poly/Bed 812. Sections for TEM were cut on a RMC PowerTomeII at 70 nm thickness stained with lead citrate and uranyl acetate. Grids were imaged on a Phillips CM200 TEM.

### Light sheet microscopy

Adult male or female *B. malayi* were individually incubated at 37 °C in RPMI 1640 with L-glutamine with 125 nM DRAQ5 (Biolegend) for 24-48 hrs, washed once in RPMI 1640 and once in ddH_2_O, then immediately transferred to 37 °C 1-1.2% low melt-temp agarose (Sigma) with 1 mM levamisole hydrochloride (≥99%, TCI America). Individuals were mounted in FEP tubing with males in 0.8 mm inner diameter (BOLA) and females in 1.6 mm inner diameter tubes (BOLA) according to published protocols.^77^ Individuals were imaged on a customized multi-view light sheet microscope similar to a previously published system.^78^ The light sheet was created using a cylindrical lens and projected into the sample via an illumination objective (Olympus #UMPLFLN10XW, 10x/0.3). The fluorescence signal was collected with another objective (Olympus #UMPLFLN20XW, 20x/0.5) perpendicular to the illumination objective. A fiber-coupled laser engine (Toptica MLE) was used as the laser source delivering excitation light at 640 nm. Images were processed into maximum intensity projections (MIPs) and stacked into time-series using Fiji^72^. Nuclei counts and location (X,Y coordinates in relation to the centrepoint of the head tip) were collected manually from individual MIPs using the multi-point tool in Fiji^72^.

### Bioinformatic analyses

Short-read RNA sequencing data was trimmed using fastp^79^ and aligned to the *B. malayi* reference genome (WormBase ParaSite^80^, release 15) using STAR^81^. The RNA-seq pipeline was implemented using Nextflow^82^ and is publicly available (https://github.com/zamanianlab/Core_RNAseq-nf). All downstream expression analyses were carried out using a mixture of custom R, bash, and Python scripts, including hierarchical clustering and visualization. Identification of robustly-expressed genes and cryosection quality control were carried out using raw gene counts, while hierarchical clustering and primary heatmap analyses were performed with scale-normalized TPM values. Differential expression analyses were carried out using DESeq2^83^. Genome-wide transmembrane prediction was performed with HMMTOP v2.1^84^. *B. malayi* cys-loop LGICs were identified using a reciprocal blastp^85^ and profile HMM^86^ approach using a database of known *C. elegans* LGICs. Ion channel subunits were aligned with MAFFT^87^ and trimmed with trimAl^88^ such that columns with greater than 30% gaps were removed, and sequences that did not have at least 70% of residues that aligned to columns supported by 70% of the sequences were removed. The trimmed, filtered alignment was subjected to maximum-likelihood phylogenetic inference with IQ-TREE 2^89^ and ModelFinder^90^ with ultrafast bootstrapping^91^, using the VT substitution matrix^92^ with empirical base frequencies and a free-rate substitution model^93^ with 10 categories. Bootstrap values from 1,000 replicates were drawn as nodal support onto the maximum-likelihood tree.

## DATA AVAILABILITY

All data and scripts used for expression analyses, comparative genomics, and data visualization are available at https://github.com/zamanianlab/BrugiaSpatial-ms. Short-read sequencing data has been deposited into NIH BioProject PRJNA548881.

## ACKNOWLEDGEMENTS

The authors thank the University of Wisconsin-Madison Biotechnology Center Gene Expression Center & DNA Sequencing Facility for providing library preparation and next generation sequencing services as well as the NIH-NIAID Filariasis Research Reagent Resource Center (via BEI Resources) for supply of *B. malayi* worms. The authors would also like to thank the members of the Zamanian laboratory for critical comments on the manuscript.

## AUTHOR CONTRIBUTIONS

M.Z. and P.A. conceived and designed the experiments. P.A. optimized spatial and tissue sequencing pipelines that were further refined and carried out by P.A., K.V., N.D., K.G., and N.W. K.V. prepared in-house RNA-seq libraries. P.A. coordinated light sheet experiments with K.S., J.He., and J.Hu., and electron microscopy with N.S. P.A. and M.Z. analyzed the primary data and wrote the paper with input and feedback from co-authors.

## MATERIALS and CORRESPONDENCE

Contact Mostafa Zamanian at:

205 Hanson Biomedical Sciences

University of Wisconsin–Madison

1656 Linden Drive

Madison, WI 53706-1581

## SUPPLEMENTARY INFORMATION

**Supplementary Table 1. Differential patterns of gene expression across head and body tissues in adult female *B. malayi*.**

**Supplementary Table 2: Clustering of *B. malayi* genes by spatial expression pattern as resolved by RNA tomography.** Gene IDs are associated with clusters shown in Figure 4F.

**Supplementary Table 3: Differential patterns of gene expression across intestinal and non-intestinal tissues in adult female *B. malayi*.**

**Supplementary Table 4: Expression patterns (TPM values) for LCM captured pharyngeal and non-pharyngeal tissues.**

**Supplementary File 1. Clade III nematode comparative anatomy references.**

**Supplementary File 2. Phylogenetic tree of *B. malayi* LGICs.** Tree includes *B. malayi* and *C. elegans* sequences.

**Supplementary Figure 1. Adult female head regions of *Brugia pahangi* and *Dirofilaria immitis* visualized by Critical Point Drying (CPD) SEM.** (A-E) *Brugia pahangi*. (A) Anterior to vulva single plane, scale = 100 μm, (B) head zoom from A, scale = 50 μm, (C) en-face view, scale = 25 μm, (D) close up of vulva, scale = 10 μm, (E) multi-focus merge from head to vulva, scale = 100 μm. (F-G) *Dirofilaria immitis* (F) head including vulva, scale = 100 μm and (G) close up of vulva, scale = 10 μm.

**Supplementary Figure 2. Spatial correlations among LGIC channel subunits.** Pearson correlation coefficients were calculated from scale-normalized TPM values for the highest-quality RNA tomography replicate.

**Supplementary Figure 3. Clustering of pharyngeal and non-pharyngeal low-input RNA-seq samples.** Samples are clustered based on euclidean distances of variance stabilizing transformed (vst) count data.

**Supplementary Video 1. Light sheet microscopy pulsing of putative ES pore and channel in live adult male *B. malayi*.** Maximum intensity projections (1 / 10s) of DRAQ5 stained nuclei in individual specimens showing anterior-most head region (scale bar 50 μm) and zoom section of pulse activity (scale bar 10 μm).

**Supplementary Video 2: Posterior-anterior flythrough of Serial Block Face-SEM.** 70 nm sections in anterior region of the adult female *B. malayi* head between the nerve ring and esophageal-intestinal boundary as viewed from the posterior end.

**Supplementary Video 3: Timelapse of adult female *B. malayi* embedding in O.C.T.** Representative methylene blue stained individual oriented perpendicular to the base mold and methylene blue dye dot, flash frozen on a bed of dry ice. Videos were used to estimate the location of the anterior tip of the parasite head relative to dye dot, and to guide a conservative cryosectioning strategy to help ensure capture of the entire parasite head region.

