## Supplementary figures and images for "Spatial transcriptomics reveals antiparasitic targets associated with essential behaviors in the human parasite *Brugia malayi*"

### Supplementary Figure 1

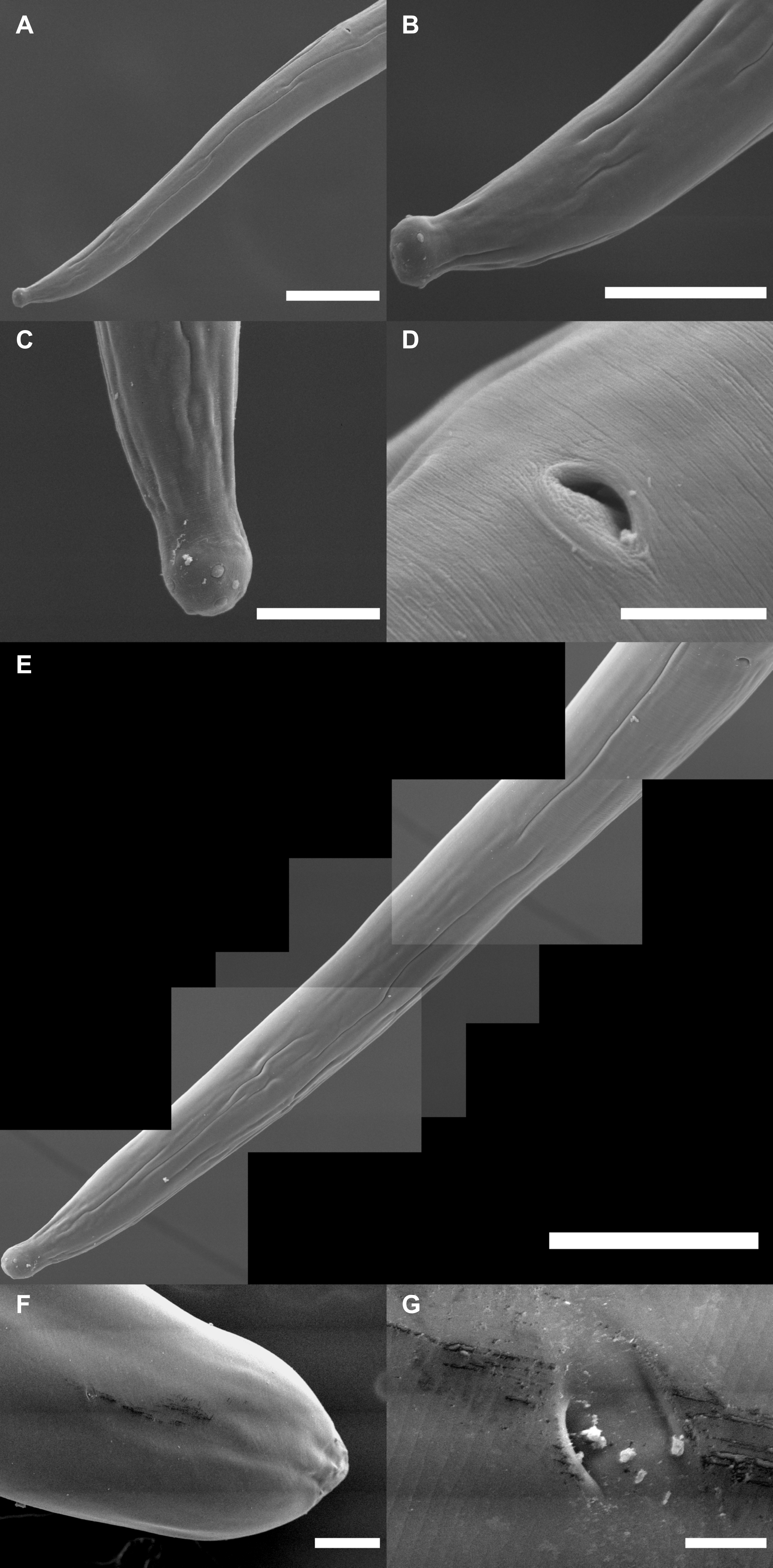

### Supplementary Figure 2

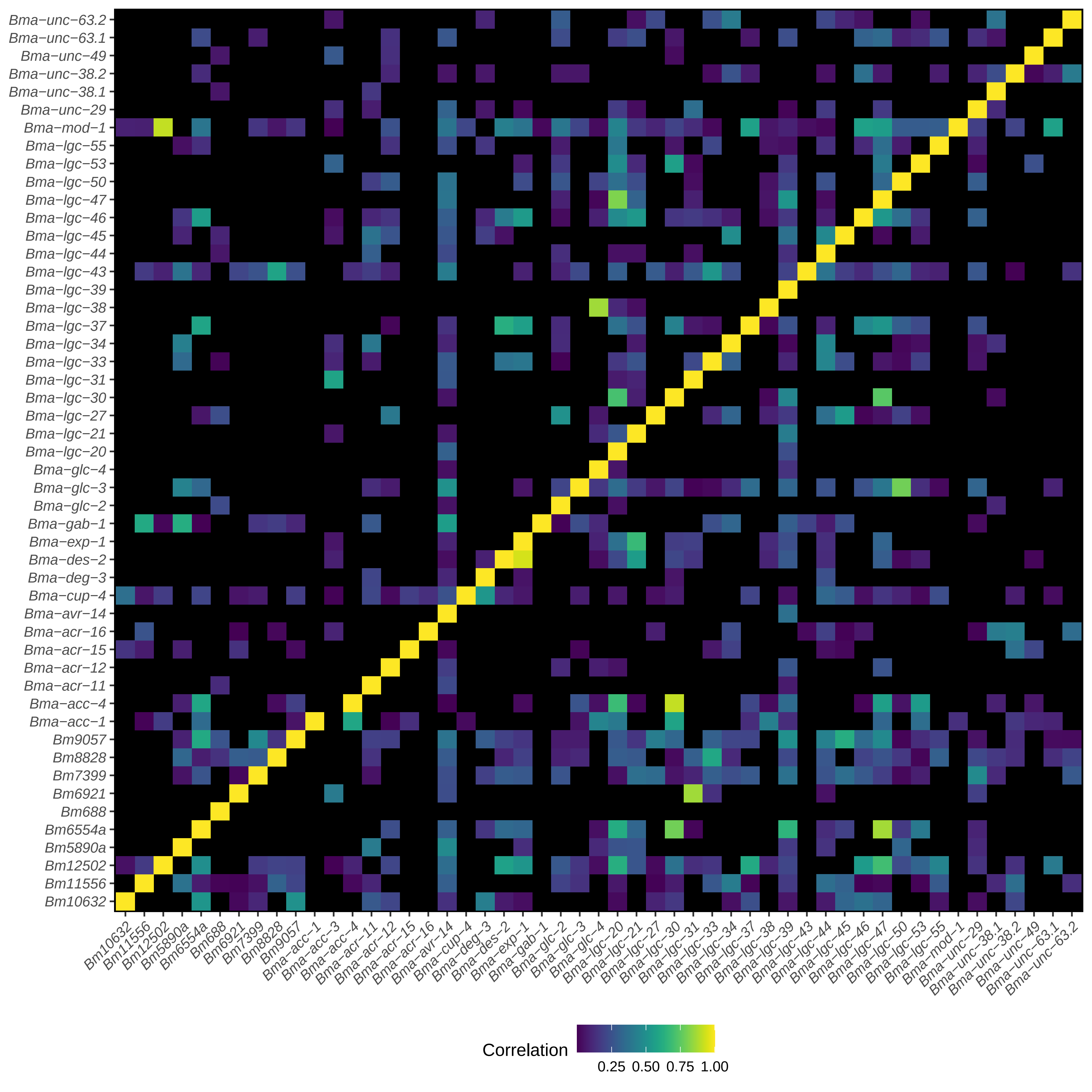

### Supplementary Figure 3

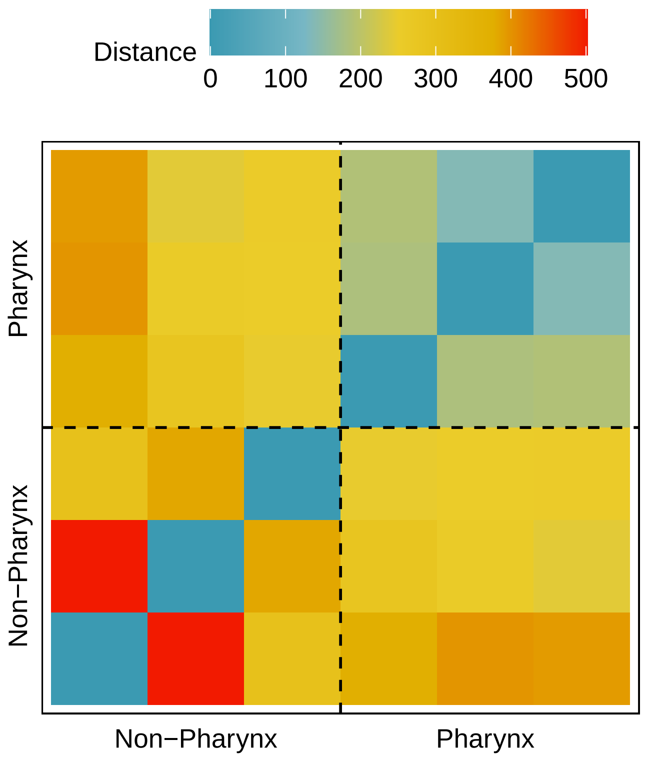
