## Supplementary File 1 for "Spatial transcriptomics reveals antiparasitic targets associated with essential behaviors in the human parasite *Brugia malayi*"

Supplementary File 1: Comparative anatomy references of Clade III nematodes

Key references denoting relative locations of key structures in Clade III nematodes selected by phylogenetic closeness to *Brugia malayi* according to previous publications.[1,2]

Species include:
*Brugia malayi [3–7], Brugia timori [8–10], Brugia pahangi [6,11–14], Wuchereria bancrofti [5,15–19], Loa loa [5,20,21], Litomosoides petteri [22], Acanthocheilonema odendhali [23–25], Elaeophora schneideri [26,27], Mansonella interstitium [28], Mansonella perstans [29], Dirofilaria immitis [5,30,31], Onchocerca volvulus [32–34], Oxyspirura conjunctivalis [35], Oxyspirura mansoni [36], Gongylonema pulchrum [37,38], Dracunculus globocephalus [39,40], Dracunculus lutrae [41], Anisakis simplex [42–44], Ascaris lumbricoides [45], Ascaris suum [46,47], Parascaris equorum [48], Toxocara canis [49–52], Enterobius anthropopitheci [53], Enterobius vermicularis [54–56],* and *Lemuricola nycticebi [57].*


1. Smythe AB, Holovachov O, Kocot KM. Improved phylogenomic sampling of free-living nematodes enhances resolution of higher-level nematode phylogeny. BMC Evol Biol. 2019;19: 121.

2. Consortium IHG, International Helminth Genomes Consortium. Comparative genomics of the major parasitic worms. doi:10.1101/236539

12. Schacher JF. Developmental stages of Brugia pahangi in the final host. J Parasitol.

1962;48: 693–706.

13. Vincent AL, Portaro JK, Ash LR. A Comparison of the Body Wall Ultrastructure of Brugia pahangi with that of Brugia malayi. J Parasitol. 1975;61: 567–570.

14. Collin WK. Ultrastructural morphology of the esophageal region of the infective larva of Brugia pahangi (nematoda: Filarioidea). J Parasitol. 1971;57: 449–468.

15. Chitwood BG, Chitwood MB, Others. An introduction to nematology. Section I. Anatomy. Monumental Printing Company, Baltimore; 1950.

28. Price DL, Others. Description of Dipetalonema interstitium n. sp. from the grey squirrel and Dipetalonema llewellyni n. sp. from the raccoon. Proc Helminthol Soc Wash.

1962;29: 77–82.

37. Alicata JE. Early Developmental Stages of Nematodes Occurring in Swine. U.S. Department of Agriculture; 1935.
